# Maternal, placental and fetal response to a non-viral, polymeric nanoparticle gene therapy in nonhuman primates

**DOI:** 10.1101/2023.06.16.545278

**Authors:** Rebecca L. Wilson, Jenna Kropp Schmidt, Baylea N. Davenport, Emily Ren, Logan T. Keding, Sarah A. Shaw, Michele L. Schotzko, Kathleen M. Antony, Heather A. Simmons, Thaddeus G. Golos, Helen N. Jones

## Abstract

**Background:** Currently, there are no placenta-targeted treatments to alter the *in utero* environment. Water-soluble polymers have a distinguished record of clinical relevance outside of pregnancy. We have demonstrated the effective delivery of polymer-based nanoparticles containing a non-viral human *insulin-like 1 growth factor* (*IGF1*) transgene to correct placental insufficiency in small animal models of fetal growth restriction (FGR). Our goal was to extend these studies to the pregnant nonhuman primate (NHP) and assess maternal, placental and fetal responses to nanoparticle-mediated *IGF1* treatment.

**Methods:** Pregnant macaques underwent ultrasound-guided intraplacental injections of nanoparticles (*GFP-* or *IGF1-*expressing plasmid under the control of the trophoblast-specific *PLAC1* promoter complexed with a HPMA-DMEAMA co-polymer) at approximately gestational day 100 (term = 165 days). Fetectomy was performed 24 h (*GFP*; n =1), 48 h (*IGF1*; n = 3) or 10 days (*IGF1*; n = 3) after nanoparticle delivery. Routine pathological assessment was performed on biopsied maternal tissues, and placental and fetal tissues. Maternal blood was analyzed for complete blood count (CBC), immunomodulatory proteins and growth factors, progesterone (P4) and estradiol (E2). Placental ERK/AKT/mTOR signaling was assessed using western blot and qPCR.

**Findings:** Fluorescent microscopy and in situ hybridization confirmed placental uptake and transgene expression in villous syncytiotrophoblast. No off-target expression was observed in maternal and fetal tissues. Histopathological assessment of the placenta recorded observations not necessarily related to the *IGF1* nanoparticle treatment. In maternal blood, CBCs, P4 and E2 remained within the normal range for pregnant macaques across the treatment period. Changes to placental ERK and AKT signaling at 48 h and 10 d after *IGF1* nanoparticle treatment indicated an upregulation in placental homeostatic mechanisms to prevent over activity in the normal pregnancy environment.

**Interpretation:** Maternal toxicity profile analysis and lack of adverse reaction to nanoparticle-mediated *IGF1* treatment, combined with changes in placental signaling to maintain homeostasis indicates no deleterious impact of treatment.

**Funding:** National Institutes of Health, and Wisconsin National Primate Research Center.

## Introduction

Fetal growth restriction (FGR) occurs in 5-10% of all pregnancies in the developed world and has higher rates in the developing world. FGR contributes significantly to perinatal morbidity and mortality, primarily due to prematurity and hypoxia ^1^. At present, there are no treatments for FGR and the childhood and adulthood sequelae comprise a significant burden on healthcare costs worldwide. Current interventions for FGR and the short- and long-term sequelae are extremely limited and commonly include iatrogenic preterm delivery and admission to the neonatal intensive care unit (NICU), a procedure that can exacerbate detrimental developmental outcomes ^2^. Therapeutic intervention during pregnancy enabling a fetus to remain *in utero* and on an upwards growth and developmental trajectory until term may have the potential to, not only avoid a premature delivery and NICU stay, but also, remove predisposition to long-term sequelae in later life such as metabolic syndrome and cardiovascular disease ^3^.

Previous studies in animal models have explored the insulin like growth factor (IGF) axis as a therapeutic target to improve fetal outcomes as it plays a major regulatory role in placental and fetal growth and is essential for normal growth and development ^4^. In many species, both *IGF1* and *IGF2* genes are expressed in fetal tissues throughout pregnancy ^5^, and reduced in FGR ^6,7^. In transgenic mouse models ^8,9^, *Igf1, Igf2* and *Igf1/Igf2* deletion results in 60% fetal weight reduction and reduced placental growth. Human studies also show that growth restricted fetuses have a reduction in IGF1, IGF2, IGFBP3 and an increase in IGFBP1 ^10,11^. Animal studies involving infusion of IGF protein into the maternal or fetal compartments have shown conflicting results. In rabbits, Skarsgard et al. ^12^ saw no increase in fetal weight following either maternal or intraamniotic delivery of IGF1, and similarly in rats, Woodall et al. ^13^ saw no change in fetal or placental weight. In contrast, Sferruzzi-Perri et al. ^14^ demonstrated in the pregnant guinea pig that chronic maternal infusion of IGF-I throughout gestation was associated with an increased placental capacity to transport glucose and amino acid transport system A activity (MeAIB) at term. However, these guinea pig studies involved long-term infusions of IGF1 starting in early to mid-gestation, a period of time before which clinical identification of FGR in human pregnancy is possible.

Water-soluble polymers have been used in clinic and/or clinical trials for the modification of proteins, modification of liposomes, surface modification of biomaterials, and as carriers of drugs, genes, and oligonucleotides ^15,16^. We have demonstrated ^17^ that poly-(2-hydroxypropyl) methacrylamide (HPMA), a non-immunogenic, water-soluble polymer is biocompatible with poly(2-(N,N-dimethylamino)) ethyl methacrylate (DMAEMA), a tertiary amine that acts as a weak base capable of being protonated at biological pH. The resulting co-polymer (HPMA-DMAEMA) can be complexed with purified plasmid DNA containing an expression cassette for a transgene (reporter or functional). The resulting plasmid-containing nanoparticles can be taken up by cells and we have demonstrated successful expression of the delivered transgene in *in vitro* and *ex vivo* human trophoblast and placenta models ^18^. We have also previously demonstrated that placental over-expression of *IGF1* in a surgically-induced mouse model of placental insufficiency can maintain appropriate fetal growth ^17^. Additionally, in the guinea pig maternal nutrient restriction (MNR) model of FGR, we demonstrated significant alterations in placental and fetal development, including increased nutrient transporter expression and normalization of gene expression changes associated with FGR in fetal tissues ^19,20^.

Rodent models of FGR, while valuable, and used for our proof-of-concept studies ^17,21,22^ are restrictive given the short gestation of the species. While guinea pigs are well suited for modeling human FGR ^23^, they are litter bearing species which presents limitations towards determining translatability of nanoparticle gene therapies. Hence, the need to use a nonhuman primate (NHP) in which to assess interventions during pregnancy is required for translation of our interventional strategies into human pregnancy with potential clinical impact. The NHP more accurately reflects human placental morphology, endocrinology, extravillous invasion and remodeling of the decidual spiral arteries, and trajectory of fetal development ^24^. The aim of the current study was to evaluate the safety of delivering placenta-specific, nanoparticle-mediated gene therapy in healthy pregnant macaques and investigate maternal, placental and fetal responses to treatment at 48 h and 10 days after delivery.

## Methods

### Animals

One pregnant cynomolgus macaque (*Macaca fascicularis*; 5.2 yrs, 4.6 kg, initial nanoparticle uptake experiment), and five pregnant rhesus macaques (*Macaca mulatta;* 4-11 yrs, 5.8-10.3 kg) were used in this study. The macaques were cared for by the staff at the Wisconsin National Primate Research Center. All procedures were performed in accordance with the NIH Guide for the Care and Use of Laboratory Animals and under the approval of the University of Wisconsin College of Letters and Sciences and Vice Chancellor Office for Research and Graduate Education Institutional Animal Care and Use Committee (animal protocol number G006040). The female macaques described in this report were co-housed with compatible males following the observation of menses. Pregnancy was detected by abdominal ultrasound and gestational age was estimated as previously described ^25^. For physical examinations, intraplacental injection, and blood collection procedures, the dams were anesthetized with an intramuscular dose of ketamine (∼10 mg/kg). Animals were monitored at least twice daily for general health and well-being. Archived placental formalin-fixed paraffin embedded blocks, flash frozen tissue and RNA from Dr. Golos’ laboratory served as non-contemporary untreated controls for immunohistochemistry, protein and RNA expression analysis, respectively.

### Intraplacental injection of nanoparticles

Synthesis of the HPMA_115_-DMAEMA_115_ co-polymer has been previously described ^19^. Nanoparticles were formed by complexing plasmids, containing either the *green fluorescent protein* (*GFP*) or *human IGF1* gene under control of the trophoblast-specific promotor *PLAC1*, with the HPMA_115_-DMAEMA_115_ co-polymer under sterile conditions. To visualize particle localization and transgene expression, Texas-red fluorophore-conjugated HPMA-DMAEMA co-polymer was complexed with PLAC1-*GFP* plasmid. All subsequent experiments were performed using an unconjugated HPMA-DMAEMA co-polymer complexed with PLAC1-*IGF1* plasmid. 150 μg of plasmid was provided in a final injection volume of 0.5 mL.

Nanoparticles were administered to the placenta by ultrasound-guided transabdominal needle placement. A 20-gauge spinal needle was advanced under ultrasound visualization through the lateral aspect of the abdomen until the tip of the needle was visualized in the placenta. The uterus and surrounding tissues were scanned to confirm lack of hemorrhage following injection. Pregnant macaques were monitored daily prior to and after placental injection for any clinical signs of impending pregnancy loss and general well-being.

### Blood and Tissue Collection

Experimental time points for maternal blood, maternal biopsies, maternal-fetal-interface and fetal tissue collection are provided in Supplemental Figure S1. Blood samples were obtained from the femoral or saphenous vein using a vacutainer system. Complete blood count tests of whole blood collected in an EDTA coated tube were performed by the WNPRC Pathology Services Unit as previously described ^26^.

At either 48 h or 10 d post-injection, the fetoplacental unit was collected via fetectomy. Maternal biopsies were collected aseptically and the dam was recovered. The fetus was euthanized by intravenous or intracardiac injection of 50 mg/kg sodium pentobarbital. A center cut of the placenta containing the umbilical cord insertion site was collected for histopathological analysis. The placental disc was then dissected into 6 regions using a grid pattern and from each grid piece a sample of tissue was preserved by flash freezing, collected into RNAlater (*ThermoFisher Scientific*), or fixed for histopathological analysis. The decidua and placenta/chorion were separated and individually collected.

### Hormone Assays

Hormone assays were performed by the Wisconsin National Primate Research Center Assays Services Unit. Maternal serum samples were analyzed for estradiol (25 μL) and progesterone (20 μL) using a Cobas e411 analyzer equipped with ElectroChemiLuminescence technology (*Roche*) according to manufacturer instructions. Results were determined via a calibration curve which was instrument-generated by 2-point calibration using traceable standards and a master curve provided via the reagent barcode. Inter-assay coefficient of variation (CV) was determined by assay of an aliquot of a pool of rhesus plasma. For estradiol, the limit of quantitation (LOQ) was 25 pg/mL, the intra-assay CV was 2.02%, and the inter-assay CV was 5.05%. For progesterone, the LOQ was 0.2 ng/mL, the intra-assay CV was 1.37%, and the inter-assay CV was 4.63%.

### Luminex Assays

Maternal peripheral blood plasma immunomodulatory proteins and growth factors were analyzed with a Cytokine/Chemokine/Growth Factor 37-plex NHP ProcartaPlex kit (*ThermoFisher Scientific*). Plasma samples were run in duplicate on a Bioplex 200 instrument (*Bio-Rad*) and were analyzed with the Bioplex Manager Software as previously described ^27^.

### Histopathology

Tissues were fixed in 4% PFA for 24 h prior to transfer to 70% ethanol. Paraffin embedding, tissue sectioning and hematoxylin and eosin (H&E) staining were performed by the University of Wisconsin Translational Research Initiatives in Pathology laboratory and the Comparative Pathology Laboratory through the University of Wisconsin Research Animal Resources and Compliance office. Histological evaluation of fetal and maternal tissues and evaluation of placenta center sections for chronic deciduitis, villitis, maternal decidual vasculitis, decidual necrosis, acute villous infarctions, and villous calcifications was done by a board-certified veterinary pathologist (HAS) blinded to the treatment.

### Fluorescent microscopy

Visualization of the nanoparticle and transgene expression was performed on samples collected from the Texas-Red conjugated, *GFP* plasmid nanoparticle treated individual. Placenta tissue was fixed in 2% PFA for 4 h, then washed and moved to 9% sucrose in PBS for 4 h, and then placed in 30% sucrose in PBS overnight at 4ºC. The tissues were stabilized in OCT and sectioned (7 μm thickness) prior to imaging. Texas Red and GFP fluorescence was visualized using the Eclipse Ti Inverted microscope (*Nikon*). Exposure time was set using untreated tissue to eliminate auto- and background fluorescence.

### In situ Hybridization

In situ hybridization (ISH) of plasmid-specific *IGF1* was performed on placental tissue collected 48 h and 10 d after the intraplacental injection of nanoparticles, and maternal and fetal liver and lung to confirm transgene expression as previously described ^18,20^. ISH was performed using a custom designed BaseScope™ probe (*Advanced Cell Diagnostics*) as outlined previously ^19^.

### Immunohistochemistry

Standard immunohistochemistry (IHC) protocols were used to confirm localization of plasmid-specific mRNA in placental trophoblast cells (cytokeratin), visualize immune cells (CD45), and trans-localization of glucose transporter SLC2A1 in 5 μm thick placenta sections. Sections were de-waxed and rehydrated as standard, and targeted antigen retrieval performed using boiling 1x Targeted Retrieval Solution (*Invitrogen*) for 20 mins. Endogenous peroxidase activity was suppressed by incubating sections in 3% hydrogen peroxide for 10 mins. Sections were then blocked in 10% goat serum with 1% bovine serum albumin (BSA; *Jackson ImmunoResearch*) in PBS for 1 h at room temperature. Anti-pan cytokeratin (*Bethyl Laboratories* A500-019A; 1:100), anti-CD45 (*Dako* M0701; 1:100) or anti-SLC2A1 (*Abcam* ab15309; 1:750) primary antibodies were applied overnight at 4ºC followed by an incubation with a biotinylated anti-mouse IgG (*Vector BA-9200*; 1:200 for CD45) or biotinylated anti-rabbit IgG (*Vector BA-1000*; 1:200 for SLC2A1) secondary antibody for 1 h at room temperature. Staining was amplified using the Vector ABC kit (*Vector*) and detected using DAB (*Vector*) for brown precipitate. Hematoxylin was used to counterstain nuclei. All sections were imaged using the Axioscan scanning microscope (*Zeiss*), and representative images were captured using the Zen Imaging software (*Zeiss*). Semi-quantitative assessment of SLC2A1 was performed by scoring staining intensity (0 = none, 1 = light, 2 = medium, 3 = dark) of the apical and basal syncytiotrophoblast membranes in 3 randomly chosen representative high-power fields of view per placenta section. Assessment was performed by 2 independent scorers, blinded to treatment.

### RNA Isolations and Quantitative PCR (qPCR)

Methods for gene expression analysis have been previously published ^19^. Briefly, approximately 50 mg of snap frozen placenta tissue was lysed in RLT-lysis buffer (Qiagen), and RNA was extracted using the RNeasy Mini kit (Qiagen) including DNase treatment following standard manufacturers protocol. 1 μg of RNA was converted to cDNA using the High-capacity cDNA Reverse Transcription kit (Applied Biosystems). For qPCR, PowerUp SYBR green kit (Applied Biosystems) was used, and included predesigned primers (*Sigma* KiQStart Primers). Gene expression was normalized using housekeeping genes *β-ACTIN, TBP* and *GAPDH*. qPCR was performed using the Quant3 Real-Time PCR System (Applied Biosystems), and relative mRNA expression calculated using the comparative CT method with the Design and Analysis Software v2.6.0 (Applied Biosystems).

### Western Blot

Methods for western blot analysis have been previously published ^19^. Briefly, placenta tissue was homogenized in ice-cold RIPA buffer containing protease and phosphatase inhibitors. 20 μg of protein was denatured by heating at 95ºC for 10 min, and then run on a 10% Tris-Bis precast gel (*Invitrogen*) following manufacturers protocols. Electrotransfer occurred onto nitrocellulose membranes using the Bolt Mini Electrophoresis unit (Invitrogen). Membranes were blocked in 5% skim-milk in Tris-buffered Saline containing Tween 20 (TBS-T) and then transferred to diluent containing primary antibodies to be incubated overnight at 4ºC. Details of primary antibodies can be found in Supplementary Table S1. The membranes were then washed and further incubated with a HRP conjugated secondary (*Cell Signaling* 7074S 1:1000) for 1.5 h at room temperature. Protein bands were visualized by chemiluminescence using SuperSignal West Femto Maximum Sensitivity Substrate (*Thermo Scientific*) on the GelDoc Imaging System (*Biorad*). Signal intensity of the protein bands were calculated using Image Lab (version 6.1, *Biorad*) and normalized to β-ACTIN or HSP60.

### Statistical Analysis

Statistical analyses were performed using SPSS Statistics 27 software. Given the small sample size, all data was assumed not to be normally distributed, and included both fetal sexes. For maternal blood measures, Related-Samples Friedman’s Two-Way ANOVA by Ranks with Bonferroni multiple testing adjustment was used to determine differences in levels 7 days prior to nanoparticle treatment, immediately before nanoparticle treatment, and at time of fetectomy. For fetal and placental weights, qPCR and western blot results, samples at 48 h and 10 d were compared to results from non-contemporary, untreated macaque weights and placenta tissue collected at a similar gestational time (n = 3-8). Statistical significance of semi-quantitative assessment of SLC2A1 staining intensity between the syncytiotrophoblast apical and basal membranes was determined using Chi-square analysis. Statistical significance was determined using generalized linear modeling including gestational day as a covariate. Results are reported as estimated marginal means ± 95% confidence interval unless otherwise stated. Statistical significance was determined at P≤0.05.

## Results

### Nanoparticle characteristics and formation

*IGF1* nanoparticles were produced using previously reported methods ^19^ under sterile conditions. Characterization of the physiochemical properties, and cellular safety and efficiency of the PHPMA_115_-b-PDMEAMA_115_ co-polymer complexed with plasmid to form the nanoparticle has been published previously ^28^.

### Placental uptake of nanoparticle and fetal response

Fetal and placental characteristics are provided in Table 1. When adjusted for gestational day, there was no difference in fetal weight at either 48 h or 10 days after *IGF1* nanoparticle treatment when compared to non-contemporary untreated controls (Table 1). Placenta disc 1 weight was lower at 48 h after *IGF1* nanoparticle treatment, as was total placental weight, when compared to untreated controls (Table 1). However, this is likely due to fetal sex. In the untreated controls, total placental weight is lower in females compared to males (estimated marginal mean weight (g) ± SEM: female = 70.9 ± 2.49 vs. male = 83.1 ± 3.37, P=0.003). Fetal sex was not determined prior to allocation of animals to these experiments, and therefore fetal sex at time of fetectomy was random. Consequently, only female fetuses were present in pregnancies collected at 48 h after *IGF1* nanoparticle treatment. Fluorescent microscopy of Texas-red and GFP fluorescence in a Cynomolgus macaque placenta confirmed nanoparticle uptake (Texas-red) and plasmid expression (GFP) 24 h after intraplacental injection (Figure 1A). Additionally, ISH for plasmid-specific *IGF1* confirmed syncytiotrophoblast uptake and plasmid expression at 48 h and, to a lesser extent, 10 days after intraplacental injection (Figure 1B & Supplemental Figure S2). Small numbers of occasional lymphocytes were observed throughout the placenta, decidua, fetal villi and fetal membranes, although this was also found in non-contemporary control specimens (Supplemental Figure S3). The overall occurrence of lesions was no different in the *IGF1* nanoparticle treated samples, compared to non-contemporary control specimens collected from normal macaque pregnancies at a similar time in gestation (Supplemental Table S2).

**Table 1.** Fetal and Placental Characteristics.

| Treatment | Dam | Fetal Sex | GD | Placenta Weight (g) |  |  | Fetal Weight (g) |
| --- | --- | --- | --- | --- | --- | --- | --- |
|  |  |  |  | Disc 1 | Disc 2 | Total |  |
| Control (Untreated) | C-1 | F | 110 | 70.0 | - | 70.0 | 175.0 |
|  | C-2 | F | 115 | 47.9 | - | 47.9 | 196.0 |
|  | C-3 | F | 107 | 47.2 | 25.8 | 73.0 | 108.5 |
|  | C-4 | F | 100 | 48.1 | 26.7 | 74.8 | 135.3 |
|  | C-5 | M | 100 | 52.0 | 29.1 | 81.1 | 148.1 |
|  | C-6 | F | 105 | 56.0 | 38.0 | 94.0 | 162.0 |
|  | C-7 | M | 100 | 52.0 | 22.0 | 74.0 | 166.0 |
|  | C-8 | M | 99 | 46.0 | 20.0 | 66.0 | 250.0 |
|  | **Mean:** |  | **103.6** | **49.9** | **26.7** | **76.2** | **165.7** |
| IGF1 48 h | IP-1 | F | 101 | 38.1 | 16.5 | 54.6 | 154.6 |
|  | IP-2 | F | 94 | 35.4 | 25.8 | 61.2 | 136.9 |
|  | IP-3 | F | 98 | 31.2 | 14.9 | 46.1 | 119.7 |
|  | **Mean:** |  | **97.7\*** | **39.8\*** | **18.4** | **57.4\*** | **176.6** |
| IGF1 10 d | IP-1 | M | 99 | 63.6 | 27 | 90.6 | 160.2 |
|  | IP-2 | M | 100 | 58.5 | 23.7 | 82.3 | 170.4 |
|  | IP-3 | F | 101 | 37.7 | 19.6 | 57.4 | 146.2 |
|  | **Mean:** |  | **100** | **52.2** | **23.8** | **75.9** | **156.9** |

Abbreviations: GD = gestational day
\*P value $\leq 0.05$ compared to Control
Estimated marginal mean based on an adjusted GD of 101

**Figure 1.**
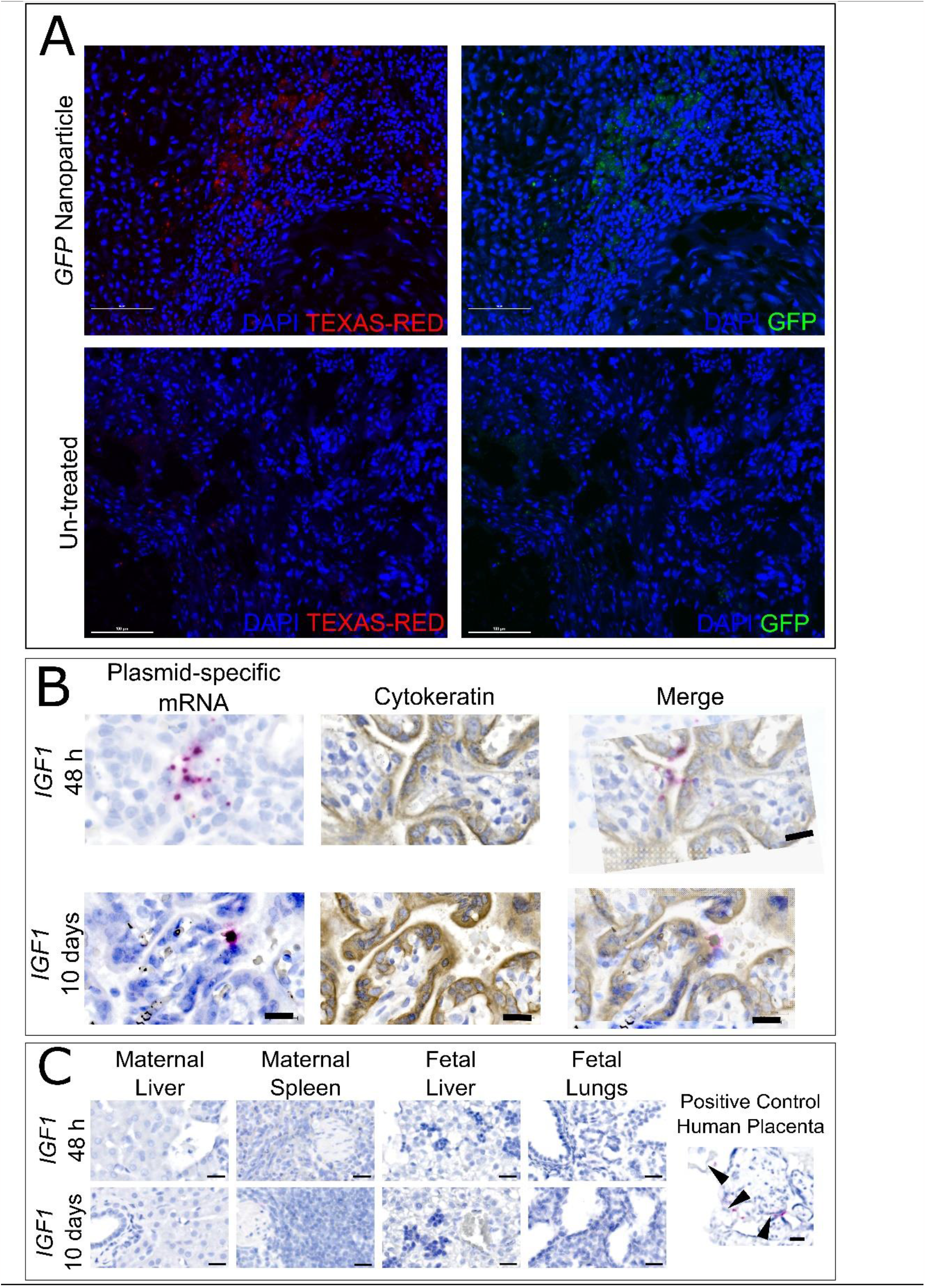
Confirmed placenta nanoparticle uptake and plasmid expression using fluorescence microscopy and *in situ* hybridization (ISH). **A**. Representative images of fluorescent microscopy for Texas-red and Green Fluorescent Protein (GFP) in macaque placenta 24 h after intraplacental *GFP* nanoparticle injection. Texas-red fluorescence confirmed nanoparticle uptake in the macaque placenta, whilst GFP fluorescence confirmed plasmid expression. Untreated macaque placenta was used as a negative control. Scale bar = 100 μm. **B**. Representative images of ISH for plasmid-specific mRNA expression in macaque placenta 48 h and 10 days after intraplacental nanoparticle treatment. ISH confirmed plasmid-specific *insulin-like 1 growth factor* (*IGF1*) expression (red dots) in the syncytiotrophoblast of macaque placenta at 48 h and 10 days after intraplacental delivery. Serial sectioning, and immunohistochemistry for the trophoblast marker cytokeratin (brown) confirmed expression in syncytiotrophoblast. Scale bar = 50 μm. **C**. Representative images of ISH for plasmid-specific mRNA expression in maternal and fetal macaque tissue after intraplacental nanoparticle treatment. ISH confirmed no off-target plasmid-specific *IGF1* expression in maternal liver nor spleen, or fetal liver and lungs, at 48 h and 10 days after intraplacental nanoparticle treatment. Human placental explant treated with nanoparticle was used as a positive control, and shows plasmid-specific *IGF1* as red dots (arrow heads) in the syncytium. Scale bar = 50 μm.

There was no indication of fetal demise following intraplacental *IGF1* nanoparticle treatment at either 48 h or 10 days. Fetal heart rate was monitored prior to and after *IGF1* nanoparticle treatment and also prior to necropsy. Fetal heart beats were within normal range, but they fluctuate depending on the level of mom’s sedation. In situ hybridization for plasmid-specific *IGF1* in fetal liver and lung at 48 h and 10 days after *IGF1* nanoparticle injection confirmed no transgene expression beyond the placenta (Figure 1C). Overall, there was no indication of severe pathology in any of the tissues at 48 h and 10 days that would indicate a negative effect on fetal development or organ function.

### Nanoparticle treatment is non-toxic in pregnant macaques

Dams were monitored at least twice daily for general well-being and no concerns were reported. At the time of fetectomy, the fetoplacental unit was collected and maternal biopsies performed on the liver, spleen, mesenteric lymph node (LN) and kidney. ISH for plasmid-specific *IGF1* expression confirmed no off-target expression in maternal liver or spleen tissue at both 48 h and 10 days (Figure 1C). Pathological assessment of maternal tissue did not indicate histopathologies beyond those observed in normal, untreated controls.

Complete blood counts (CBC) were evaluated in whole blood collected 7 days prior to *IGF1* nanoparticle injection, immediately before *IGF1* nanoparticle injection and at the time of fetectomy (48 h or 10 days) (Supplemental Figure S4). Prior to *IGF1* nanoparticle injection, the majority of the CBC parameters were within normal range, and remained within the normal range immediately before injection and at the time of fetectomy. Maternal peripheral blood immunomodulatory proteins and growth factors were analyzed and are presented in Table 2. There were minimal changes to immunomodulatory proteins/growth factors at time of *IGF1* nanoparticle injection or fetectomy, compared to pre-injection. BLC (CXCL13) was lower at 10 days after injection compared to pre-injection, whilst SDFα was higher at 48 h after injection compared to pre-injection. Finally, compared to prior to injection, progesterone and estradiol levels remained unchanged at time of intraplacental *IGF1* treatment (Table 2). There was a decrease in progesterone levels at 10 days after intraplacental *IGF1* treatment, however levels remained within the normal range for pregnant macaques ^29^.

**Table 2.** Maternal peripheral blood immunomodulatory proteins and growth factors.

| Analyte | Mean (pg/mL)<br>(95% confidence interval) |  |  |  | P<br>value |
| --- | --- | --- | --- | --- | --- |
|  | Pre (-7 days) | 0 days | 48 h | 10 days |  |
| BLC<br>(CXCL13) | 47.89<sup>a</sup><br>(30.55-75.06) | 38.96<sup>a,b</sup><br>(24.85-61.06) | 41.06<sup>a,b</sup><br>(26.20-64.37) | 37.07<sup>b</sup><br>(19.63-70.01) | **0.036** |
| Eotaxin<br>(CCL11) | 62.10<sup>a,b</sup><br>(43.44-88.78) | 50.97<sup>a,b</sup><br>(35.65-72.86) | 75.27<sup>a</sup><br>(52.65-107.60) | 36.16<sup>b</sup><br>(21.81-59.94) | **0.042** |
| ITAC<br>(CXCL11) | 11.98<br>(5.21-27.53) | 11.28<br>(4.91-25.91) | 9.69<br>(4.22-22.26) | 16.33<br>(5.03-52.94) | 0.801 |
| IL12p70 | 0.74<br>(0.57-0.95) | 0.57<br>(0.43-0.72) | 0.55<br>(0.43-0.71) | 0.50<br>(0.35-0.71) | 0.707 |
| IL1RA | 109.62<br>(61.58-195.15) | 109.03<br>(61.24-194.08) | 148.16<br>(83.22-263.75) | 148.74<br>(65.80-336.23) | 0.457 |
| IL4 | 4.66<br>(2.20-9.87) | 3.46<br>(1.38-8.68) | 2.78<br>(1.22-6.34) | 0.69<br>(0.117-4.33) | 0.392 |
| IL8 | 73.04<br>(34.53-154.49) | 46.52<br>(22.01-98.47) | 39.20<br>(18.53-82.91) | 23.54<br>(8.95-61.93) | 0.896 |
| MCP1<br>(CCL2) | 104.73<br>(76.76-142.87) | 103.02<br>(75.51-140.54) | 89.65<br>(65.71-122.30) | 80.60<br>(51.95-125.05) | 0.284 |
| MIP1α | 4.25<br>(2.72-6.66) | 3.91<br>(2.49-6.13) | 4.11<br>(2.62-6.43) | 3.75<br>(1.99-7.08) | 0.972 |
| MIP1β | 11.74<br>(6.10-22.61) | 18.82<br>(9.05-39.15) | 19.30<br>(10.61-35.11) | 12.14<br>(5.21-28.28) | 0.392 |
| NGFβ | 16.97<br>(7.83-36.79) | 10.56<br>(4.87-22.90) | 6.77<br>(2.85-16.07) | 3.03<br>(0.89-10.27) | 0.145 |
| PDGFBB | 42.93<br>(28.88-63.80) | 27.47<br>(18.49-40.83) | 52.41<br>(35.26-77.89) | 21.43<br>(12.24-37.53) | 0.468 |
| SCF | 5.59<br>(4.19-7.45) | 5.433<br>(4.07-7.24) | 5.59<br>(4.19-7.46) | 3.88<br>(2.58-5.82) | 0.457 |
| SDF1α<br>(CXCL12) | 174.08<sup>a</sup><br>(112.50-269.37) | 175.76<sup>a,b</sup><br>(113.59-271.97) | 235.81<sup>b</sup><br>(152.40-364.88) | 180.92<sup>a,b</sup><br>(97.58-335.42) | **0.042** |
| VEGFD | 45.11<br>(27.13-75.02) | 47.67<br>(28.67-79.26) | 40.68<br>(24.47-67.64) | 38.28<br>(18.65-78.57) | 0.615 |
| Progesterone | 3716.7<sup>a</sup><br>(2715.7-5086.5) | 3058.3<sup>a,b</sup><br>(2234.7-4185.6) | 3588.3<sup>a</sup><br>(2621.9-4910.9) | 2226.7<sup>b</sup><br>(1428.7-3470.3) | **0.042** |
| Estradiol | 367.02<br>(281.99-477.68) | 424.55<br>(326.20-552.56) | 479.45<br>(368.38-624.01) | 531.97<br>(366.46-772.22) | 0.978 |

P values calculated using a Related-Samples Friedman's Two-Way Analysis of Variance by Ranks. Different letters within a row denote a significant difference of $P < 0.05$ , adjusted by the Bonferroni correction for multiple tests. n = 6 dams (Pre, 0 days and 48 h) and 3 dams (10 days)

### Signaling effects of *IGF1* nanoparticle treatment in macaque placenta

Compared to untreated, there was no difference in placental expression of total AKT and total ERK at 48 h and 10 d (Figure 2A and Supplemental Figure S5). Expression of phospho-AKT did not change with *IGF1* nanoparticle treatment (Supplement 5). However, expression of phospho-ERK, and its downstream effector phospho-p90RSK was increased at 48 h and 10 d after *IGF1* nanoparticle treatment compared to untreated (Figure 2A). Additionally, expression of the mTORC1 inhibitor DEPTOR was increased, whilst expression of RAPTOR was decreased at 48 h after *IGF1* nanoparticle treatment compared to untreated (Figure 2A). At 10 d after *IGF1* nanoparticle treatment, expression of DEPTOR and RAPTOR was no different to untreated. There was also no difference in the expression of mTOR, MIOS, phospho-MEK, WDR59, NFkB, or IkB with *IGF1* nanoparticle treatment when compared to untreated (Supplemental Figure S6).

**Figure 2.**
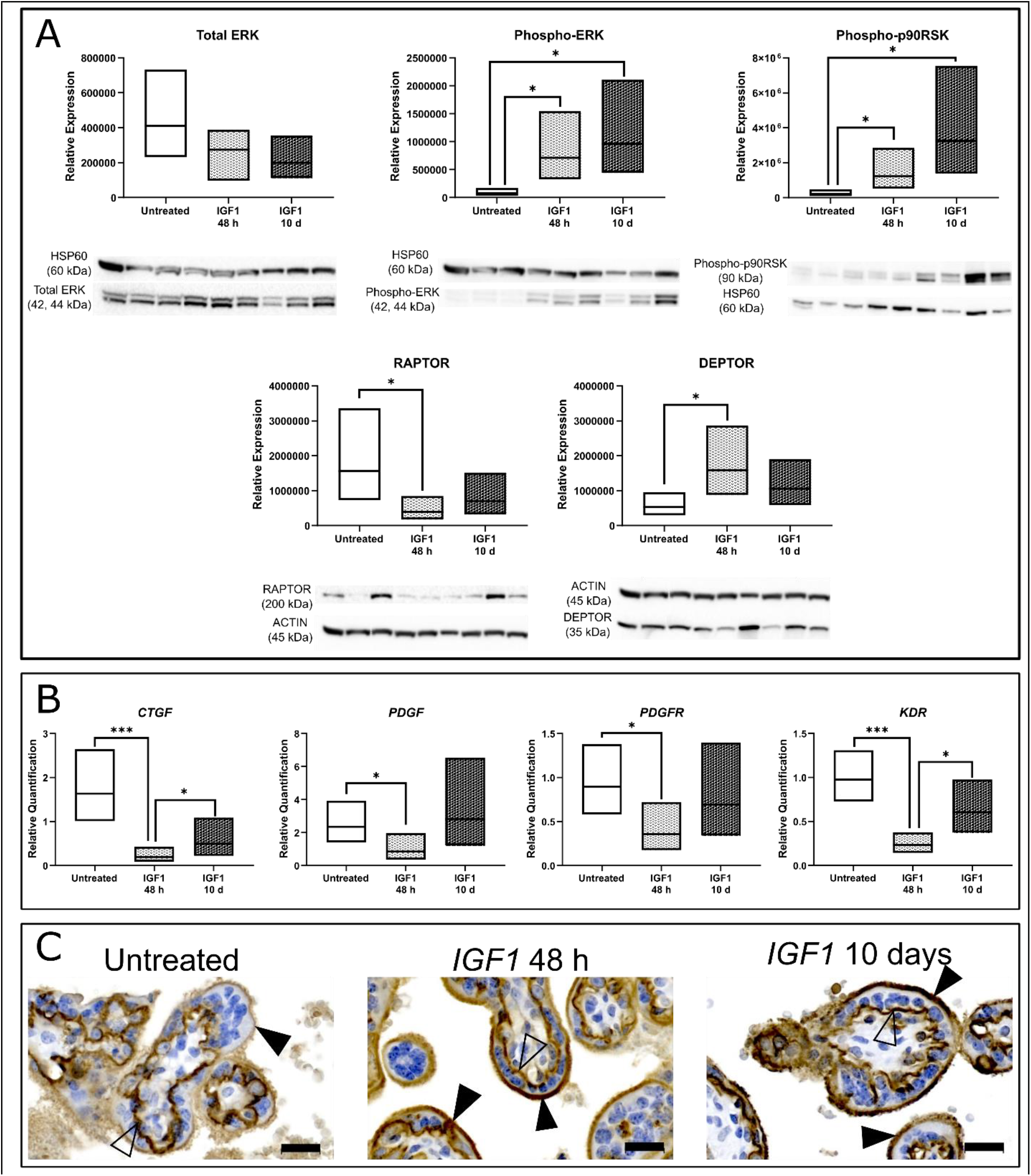
Signaling effects of *insulin-like 1 growth factor* (*IGF1*) nanoparticle treatment in macaque placenta. **A**. Protein expression of ERK and mTOR signaling members in macaque placentas 48 h and 10 d after *IGF1* nanoparticle injection. There was no difference in total ERK expression in the placenta 48 h or 10 d following *IGF1* nanoparticle, however, expression of phospho-ERK and phospho-90RSK was increased at 48 h and 10 d when compared to non-contemporary untreated controls. At 48 h after *IGF1* nanoparticle treatment, expression of RAPTOR was decreased and DEPTOR increased compared to untreated controls. There was no difference in RAPTOR or DEPTOR expression between untreated controls and 10 d after *IGF1* nanoparticle treatment. Western blot images for each protein are presented below the corresponding graph. Sample left-right: Untreated (3), *IGF1* 48 h (3), *IGF1* 10 d (3). n = 3 placentas per group. **B**. mRNA expression of growth factors in macaque placentas 48 h and 10 d after *IGF1* nanoparticle injection. Expression of *Cytokine Growth Factor* (*CTGF*) and *Vascular Endothelial Growth Factor Receptor 2* (*KDR*) was decreased in the placenta 48 h following *IGF1* nanoparticle treatment when compared to non-contemporary untreated controls and placentas collected 10 d after *IGF1* nanoparticle treatment. Expression of *Platelet Derived Growth Factor* (*PDGF*) and *PDGF Receptor* (*PDGFR*) was decreased in the placenta 48 h following *IGF1* nanoparticle treatment, but not different at 10 d after *IGF1* nanoparticle treatment when compared to non-contemporary untreated controls. n = 8 placentas untreated placentas and 3 placentas per *IGF1* nanoparticle treated group. **C**. Representative immunohistochemistry images of Glucose Transporter SLC2A1 in macaque placenta 48 h and 10 d after *IGF1* nanoparticle injection. Comparing SLC2A1 staining intensity in the syncytiotrophoblast apical (closed arrow) and basal (open arrow) membranes, in non-contemporary untreated controls, intensity was greater in the basal membrane. At 48 h and 10 d following placental *IGF1* nanoparticle treatment, staining intensity of SLC2A1 was similar between the basal and apical membranes. n = 3 placentas per group. Scale bar = 50 μm. Data are shown as the mean ± 95% confidence interval. P values calculated using a Generalized linear model and Bonferroni post-hoc analysis. *P<0.05

mRNA expression of growth factors *CTGF, PDGF, PDGF Receptor* (*PDGFR*) and *VEGF Receptor 2* (*KDR*), which are downstream effectors of AKT/ERK signaling, were decreased 48 h after *IGF1* nanoparticle treatment compared to untreated (Figure 2B). At 10 d after *IGF1* nanoparticle treatment, there was no difference in mRNA expression of *CTGF, PDGF, PDGFR* and *KDR* compared to untreated (Figure 2B). There was no difference in mRNA expression of *VEGFA, VEGFB, VEGFR1* or *PGF* at 48 h and 10 d after *IGF1* nanoparticle treatment when compared to untreated (data not shown).

Finally, in non-contemporary untreated control placenta tissue, staining intensity for glucose transporter SLC2A1 was greater in the syncytiotrophoblast basal membrane compared to the apical membrane (χ^2^ = 16.00, P<0.001; Figure 2C and Supplemental Figure S6). At 48 h and 10 d after *IGF1* nanoparticle treatment, staining intensity of SLC2A1 was comparable between the syncytiotrophoblast basal and apical membranes (χ^2^ = 0.23, P>0.05 and χ^2^ = 0.00, P>0.05, respectively), indicating re-localization to the apical surface of the fetal villi.

## Discussion

Here, we show safe non-toxic and efficient delivery of nanoparticle-mediated placenta-expressing plasmid following ultrasound-guided intraplacental injection in pregnant macaques. We observed no off-target expression in maternal or fetal tissues, and no adverse reactions to intraplacental nanoparticle treatment, similar to outcomes previously reported in mice and guinea pigs ^17,19,30^. Assessment of placental tissue demonstrated the presence of histopathological lesions at similar rates found in control, untreated macaques. Maternal complete blood counts (CBCs), cytokines and peripheral blood immunomodulatory proteins, and circulating pregnancy hormones progesterone and estradiol levels remained within normal ranges across the experimental period, suggesting the treatment was non-toxic. With only slight changes observed due to intraplacental *IGF1* nanoparticle treatment. The lack of widespread adverse acute reaction to intraplacental *IGF1* nanoparticle treatment, combined with outcomes already determined in mouse and guinea pig models, provides the necessary platform for further development and evaluation of the safety of longer-term treatment, and potential translation into the clinic as a therapeutic strategy to treat major obstetric complications.

Major obstetric complications, including preeclampsia and fetal growth restriction (FGR) contribute to fetal demise, stillbirth, NICU admission and lifelong morbidity and mortality in infants and mothers ^31^. Additionally, pregnancy complications transcend pregnancy and impact the child’s health well beyond the perinatal period and across the lifespan. Babies born from pregnancies complicated by preeclampsia, FGR and preterm birth are at increased risk of developing non-communicable diseases (NCDs) including, cardiovascular disease, hypertension, central obesity, type 2 diabetes mellitus and respiratory disease ^3,32,33^. NCDs are responsible for 80% of adult deaths annually worldwide ^34^, and have the greatest impact on health adjusted life expectancy and quality of life ^35^. “Developmental Origins of Health and Disease” (DoHAD) hypothesizes that early environmental stressors during critical fetal developmental windows result in permanent adaptive structural and physiologic changes, called developmental programming, that predispose the offspring to metabolic, endocrine, and cardiovascular diseases in postnatal life ^36^. Placental dysfunction contributes significantly to the environmental stress linked to developmental programming ^37^. Hence, the placental is the ideal site for a targeted intervention as restoration or improvement in placental function will positively impact fetal growth and development, and there are no potential long-term consequences for maternal health as the placenta is discarded after birth.

Our prior in vivo studies across multiple species/models have supported the use of a non-viral, polymer-based nanoparticle that facilitates transient gene delivery specifically to placental trophoblast cells as presented here ^17-20,30,38,39^. Similar to outcomes presented in the macaque, these studies proved the placental nanoparticle gene therapy to be safe to both mother and fetus with no observed increased fetal loss, necrosis/immune cell infiltration in the placenta, nor adverse maternal reactions. We have consistently shown no off-target nanoparticle uptake or plasmid expression in fetal tissues including brain, liver and lungs ^19,20^, and increased fetal blood glucose has been demonstrated following placental *IGF1* nanoparticle treatment in a guinea pig model of FGR ^19^, indicating the ability to modify developmental programming in fetal organs and tissues by increasing a critical resource required for adequate fetal growth. As presented here, robust histopathological assessment of maternal, placental and fetal tissues recorded no observations above normal variation in control samples. Additionally, there were very few differences in maternal blood circulating factors after *IGF1* nanoparticle treatment when compared to samples collected 7 days prior to administration.

The safety of the *IGF1* nanoparticle is further supported by minimal maternal immune response to placental administration. Assessment of cytokine expression in maternal circulation revealed very few changes following *IGF1* nanoparticle treatment when compared to expression 7 days prior to treatment. Only two were significantly different. BCL (CXCL13) was reduced in maternal circulation at 10 d after *IGF1* nanoparticle treatment compared to 7 days prior to treatment. BCL is a potent chemokine that attracts B and T lymphocytes to areas of inflammation and infection ^40,41^. In human pregnancy, BCL is present in serum and amniotic fluid of normal pregnancies and drops with advancing gestation ^42^. Hence, reduced expression of BCL at 10 d after *IGF1* nanoparticle treatment may represent the normal gestational decrease. Expression of SDF1 (CXCL12) was increased in maternal serum at 48 h following *IGF1* nanoparticle treatment compared to 7 days prior to treatment. SDF1 is a chemokine expressed in all trophoblast subtypes of the placenta from first trimester until term and involved in the modulation of placental angiogenesis ^43^. Therefore, increased expression of SDF1 48 h after IGF1 nanoparticle treatment may represent an acute change in placental expression of SDF1 related to changes in placental signaling by increased *IGF1* expression, although levels returned to baseline 10 d after *IGF1* nanoparticle treatment.

IGF1 modulates trophoblast biology through numerous different pathways including ERK and AKT signaling pathways ^44^. In situ hybridization confirmed expression of plasmid-specific *IGF1* in the macaque placenta at 48 h after *IGF1* nanoparticle administration, and to a lesser extent at 10 d after *IGF1* nanoparticle administration. Increased *IGF1* expression resulted in increased ERK signaling activity through increased expression of phospho-ERK and phospho-p90RSK. Activation of p90RSK promotes the phosphorylation of RAPTOR leading to the stimulation of mTORC1 signaling ^45^. Whilst this scenario would be appropriate in situations of placental dysfunction, in the present study, placental function was normal. Increased *IGF1* also resulted in reduced expression of RAPTOR and increased expression of the mTOR inhibitor DEPTOR at 48 h after IGF1 nanoparticle treatment, indicating a shift in placental homeostatic mechanisms to prevent over activity. RAPTOR is a scaffolding protein which recruits mTOR substrates, resulting in the regulation of mRNA translation and ribosome biogenesis ^46,47^. DEPTOR on the other hand, is the only protein reported to bind and inhibit mTOR complexes ^48^. Reduced mRNA expression of downstream mTOR effectors (i.e., *CTGF, PDGF, PDGFR* and *KDR*) further corroborates our hypothesis that under normal pregnancy environments, the placenta responds to maintain homeostasis. Additionally, it further solidifies the safety of this treatment for clinical applications, whereby misdiagnosis of placental dysfunction may occur but would not result in excess placental function with trophoblast-specific nanoparticle-mediated IGF1 delivery.

There are several limitations to the current study, including the small sample size. Given the small sample size, sex-specific differences in placental response to nanoparticle treatment could not be independently explored. However, future studies are planned to expand upon these investigations. A relatively short window of observation (up to 10 days after intraplacental *IGF1* nanoparticle treatment), which cannot exclude longer-term effects on maternal and fetal health and the single nanoparticle dose, requires additional preclinical studies designed to evaluate safety, efficacy and immunogenicity of multiple treatments. The findings presented here expand on the safety and efficacy of nanoparticle-mediated plasmid deliveryin the treatment of placenta dysfunction shown to be effective in mice and guinea pigs ^17,19,30^, further alleviating concerns for potential toxicity to mother or fetus. Overall, such information will contribute to paving the way to future clinical testing with the aim of broadening applicability of placental gene therapy to treat major obstetric complications, and possibly reduce the risk of other diseases like cardiovascular disease, type 2 diabetes and cancer in later life.

## Acknowledgments

We would like to thank Drs Craig Duvall and Mukesh Gupta at Vanderbilt University for providing the co-polymer. We also thank the Wisconsin National Primate Research Center (WNPRC) Veterinary Services, Scientific Protocol Implementation, Pathology Services, Assay Services and Animal Services Unit staff for providing animal care, and assisting in procedures including breeding, pregnancy monitoring and sample collection.

## Ethics Statement

All procedures were performed in accordance with the NIH Guide for the Care and Use of Laboratory Animals and under the approval of the University of Wisconsin College of Letters and Sciences and Vice Chancellor Office for Research and Graduate Education Institutional Animal Care and Use Committee (animal protocol number G006040).

## Data Availability Statement

The original contributions presented in the study are included in the article/Supplementary Materials, further inquiries can be directed to the corresponding author.

## Author Contributions

RLW performed experiments, analyzed data and wrote manuscript. JKS designed, oversaw experiments and contributed to writing of the manuscript. BND performed experiments and analyzed data. ER performed experiments and analyzed data. LTK performed experiments, analyzed data, organized sample collection and processed samples. SAS organized sample collection and processed samples. MLS performed animal procedures and collected tissues. KMA performed animal procedures. HAS performed necropsies and histopathological analysis. TGG designed and oversaw experiments. HNJ obtained funding, conceived the study and edited the manuscript. All authors have approved the final version.

