## Supplemental Material for "Maternal, placental and fetal response to a non-viral, polymeric nanoparticle gene therapy in nonhuman primates"

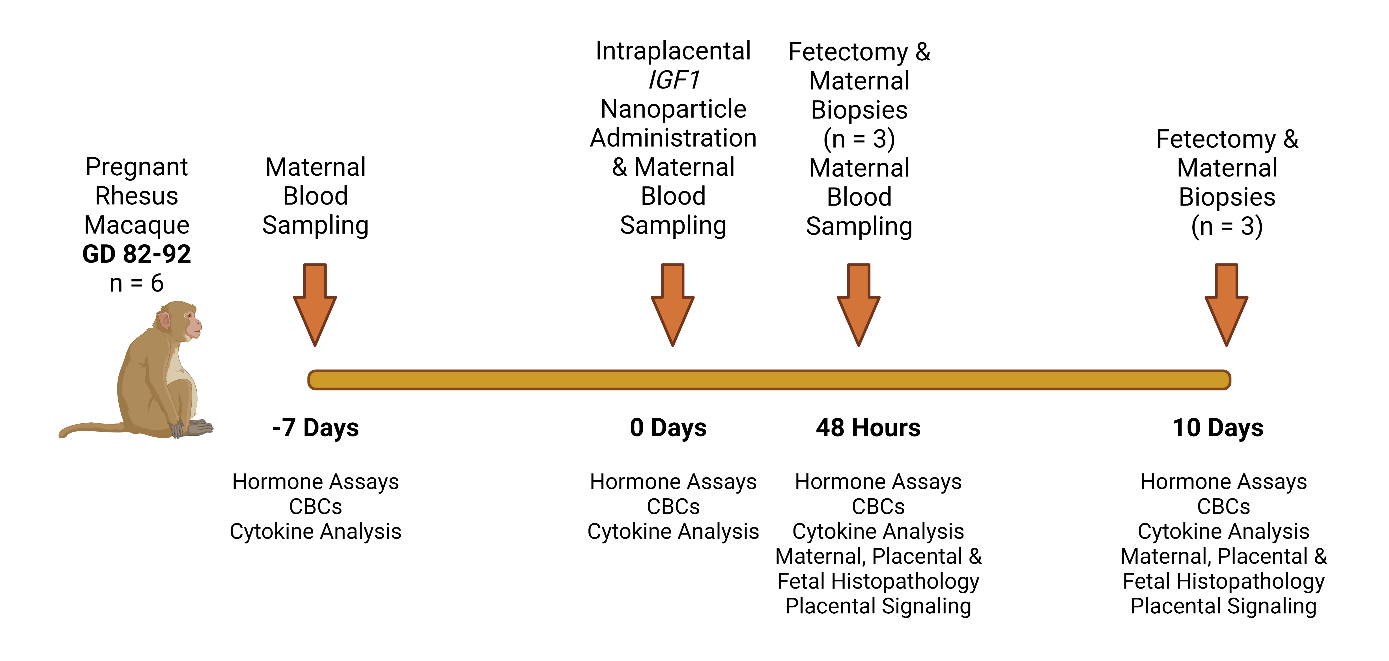

**Supplemental Figure S1**. **Experimental timepoints for maternal blood, maternal biopsies, maternal-fetal-interface and fetal tissue collection**

| Supplemental Table S1. Antibodies used for Western Blot Analysis of Placenta Signaling | | | |
| --- | --- | --- | --- |
| Antigen | **Manufacturer** | **Catalog Number** | **Dilution** |
| Total AKT | Cell Signaling | 9272 | 1:1000 |
| Phospho-AKT (S473) Pan Specific | R&D Systems | AF887 | 1:500 |
| p44/42 MAPK (ERK1/2) (137F5) | Cell Signaling | 4695 | 1:1000 |
| Phospho-p44/42 MAPK (ERK1/2) (Thr202/Tyr204) | Cell Signaling | 4370 | 1:2000 |
| Phospho-p90RSK (Ser380) | Cell Signaling | 11989 | 1:1000 |
| RAPTOR (24C12) | Cell Signaling | 2280S | 1:1000 |
| DEPTOR (DEPDC6) | LSBio | LS-C187268 | 1:1000 |
| Phospho-MEK1/2 (Ser217/221) | Cell Signaling | 9154 | 1:1000 |
| mTOR (7C10) | Cell Signaling | 2983 | 1:1000 |
| MIOS (D12C6) | Cell Signaling | 13557 | 1:1000 |
| WDR59 (D4Z7A) | Cell Signaling | 53385 | 1:1000 |
| RICTOR (53A2) | Cell Signaling | 2114S | 1:1000 |

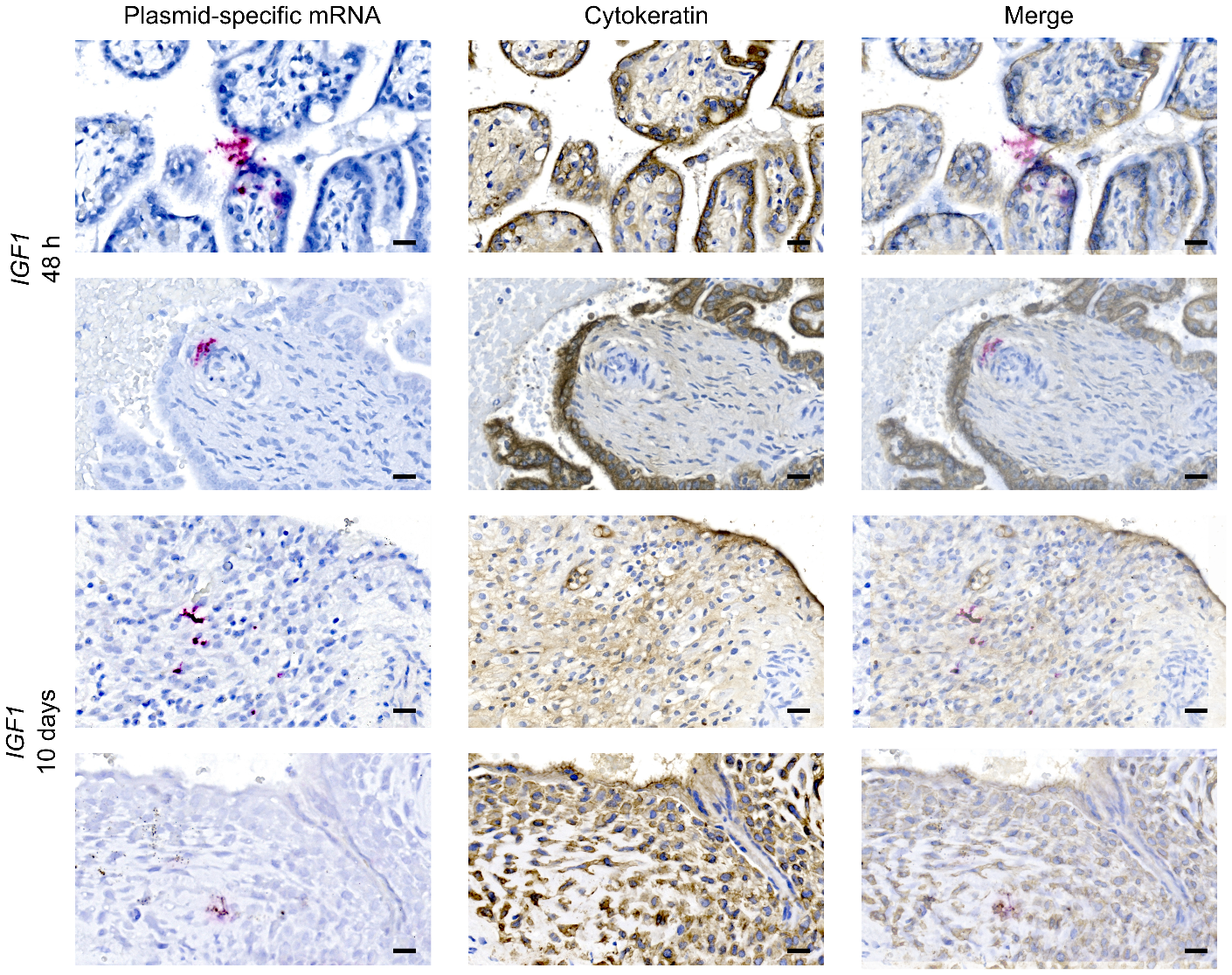

**Supplemental Figure S2**. **Additional representative images of situ hybridization (ISH) for plasmid-specific mRNA expression in macaque placenta 48 h and 10 days after intraplacental nanoparticle treatment.** Serial sectioning, and immunohistochemistry for the trophoblast marker cytokeratin (brown) confirmed expression in syncytiotrophoblast. Scale bar = 20 µm.

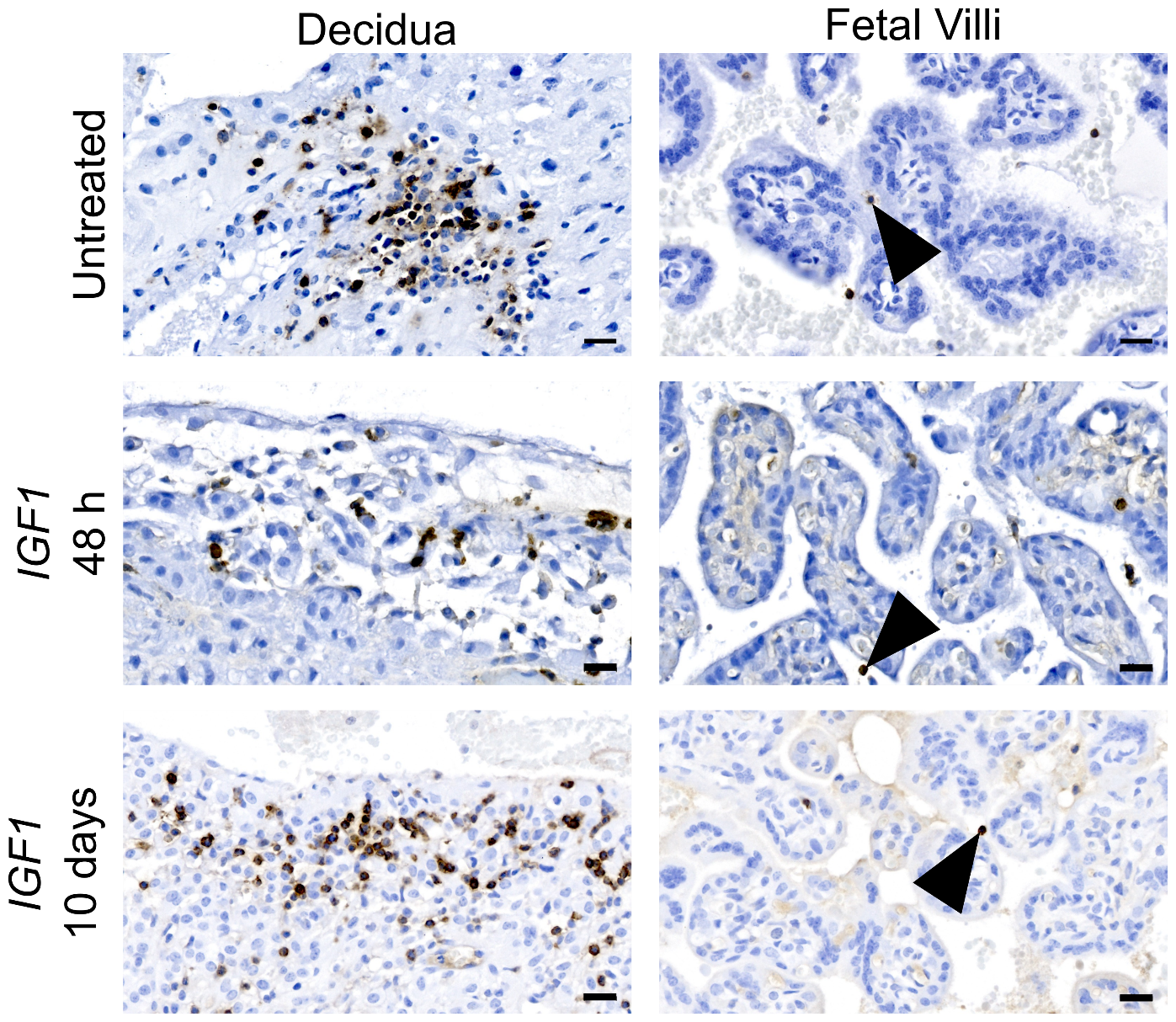

**Supplemental Figure S3. Representative images of immune cells in the macaque decidua and fetal villi.** Immunohistochemistry was used to identify CD45 positive immune cells in the decidua and fetal villi of untreated placenta tissues, and placenta tissue 48 h and 10 days following intraplacental nanoparticle treatment. N = 3 macaques per group. Scale bar = 20 µm

| **Supplemental Table S2**. Overall occurrence of lesions in macaque placenta 48 h and 10 days after intraplacental *IGF1* nanoparticle treatment compared to non-contemporary, untreated control specimens | | | |
| --- | --- | --- | --- |
|  | **Untreated (control)** | **48 h** | **10 days** |
| Chronic Deciduitis | 5/8 | 1/3 | 1/3 |
| Villitis | 3/8 | 2/3 | 2/3 |
| Maternal Decidual Vasculitis | 2/8 | 2/3 | 1/3 |
| Acute Villous Infarction | 3/8 | 2/3 | 1/3 |
| Villous Mineralization | 7/8 | 3/3 | 2/3 |
| Decidua Necrosis | 2/8 | 0/3 | 1/3 |
| Placental Necrosis | 6/8 | 3/3 | 3/3 |
| Hemorrhage | 6/8 | 3/3 | 3/3 |
| Ischemia | 1/8 | 2/3 | 1/3 |
| Acute Intervillositis | 4/8 | 1/3 | 2/3 |
| Villous Fibrin Accumulation | 3/8 | 2/3 | 3/3 |
| Hemosiderosis/Hemosiderin accumulation | 4/8 | 3/3 | 2/3 |

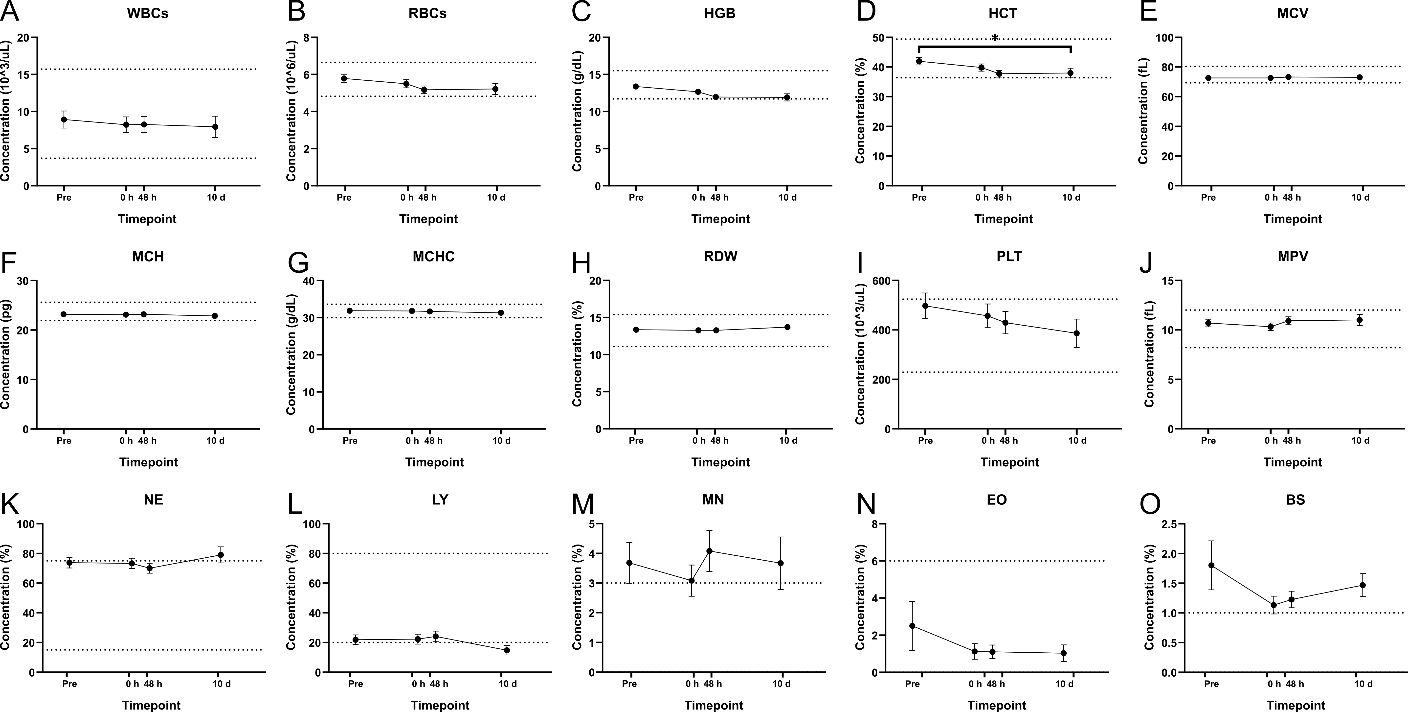

**Supplemental Figure S4 Complete blood counts (CBC) in maternal macaque whole blood prior to *IGF1* nanoparticle injection (Pre), immediately before *IGF1* nanoparticle injection (0 h) and at the time of fetectomy (48 h and 10 d).** There was no change to maternal whole blood CBC measures due to intraplacental *IGF1* nanoparticle injection. **A** WBCs: white blood cells. **B** RBCs: red blood cells. **C** HGB: hemoglobin. **E** MCV: mean corpuscular volume. **F** MCH: mean corpuscular hemoglobin. **G** MCHC: mean corpuscular hemoglobin concentration. **H** RDW: red blood cell distribution width. **I** PLT: platelets. **J** MPV: mean platelet volume. **K** NE: neutrophils. **L** LY: lymphocytes. **M** MN: monocytes. **N** EO: eosinophils. **O** BS: basophils. n = 6 dams (Pre, 0 h and 48 h) and 3 dams (10 d). Data are mean ± SEM. P values calculated using a Related-Samples Friedman's Two-Way Analysis of Variance by Ranks. *P<0.05

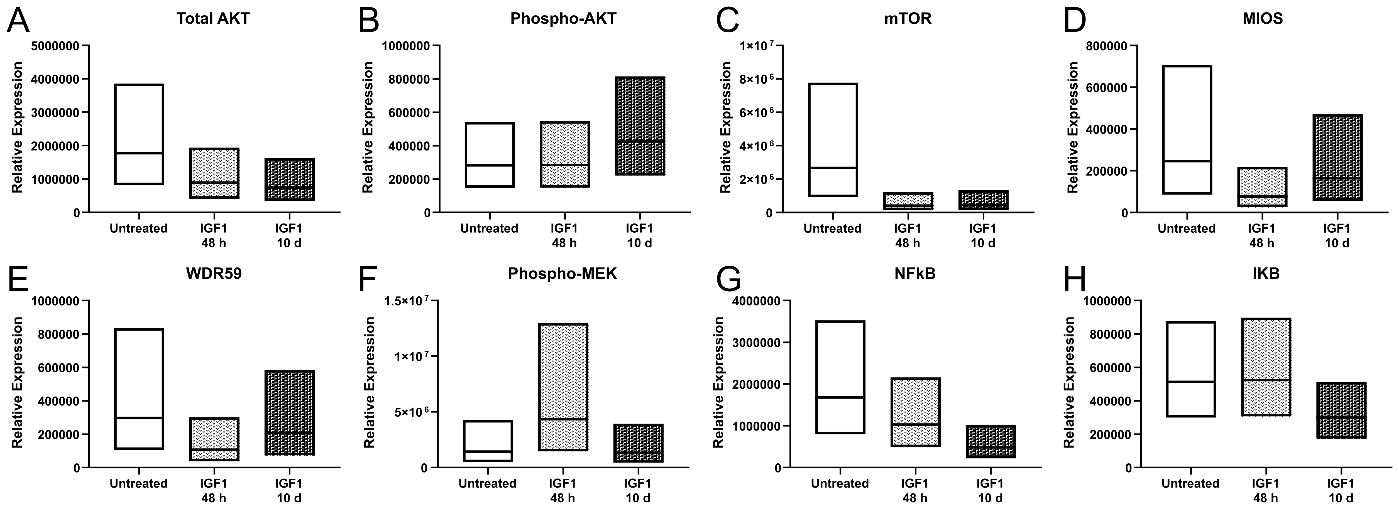

**Supplemental Figure S5. Protein expression of ERK and mTOR signaling members in macaque placentas 48 h and 10 d after *IGF1* nanoparticle injection.** There was no difference in total AKT (**A**), Phospho-AKT (**B**), mTOR (**C**), MIOS (**D**), WDR59 (**E**), Phospho-MEK (**F**), NFkB (**G**) or IkB (**H**) expression in the placenta 48 h or 10 d following *IGF1* nanoparticle treatment when compared to non-contemporary untreated controls. n = 3 placentas per group. Data are mean ± 95% confidence interval.

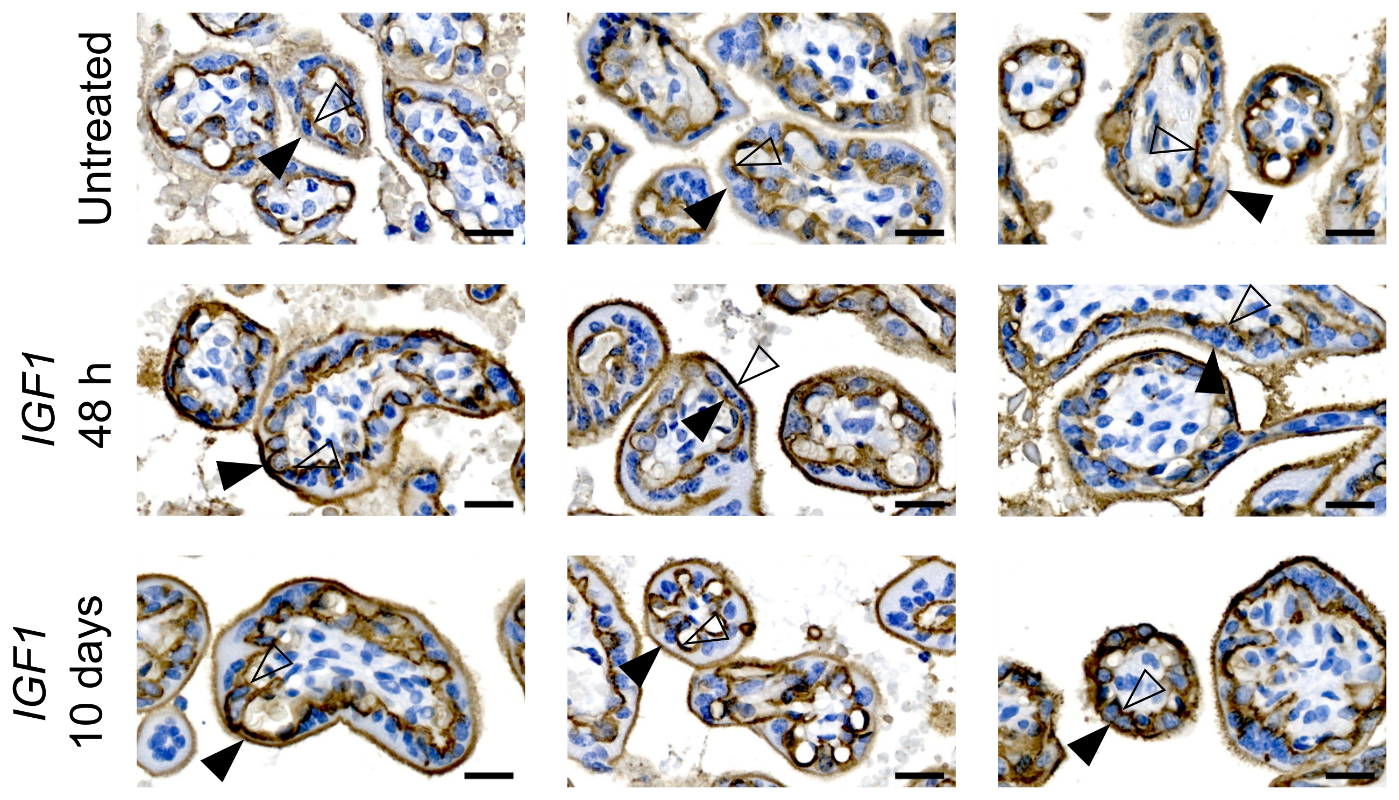

**Supplemental Figure S6. Representative immunohistochemistry images of Glucose Transporter SLC2A1 in macaque placenta 48 h and 10 d after *IGF1* nanoparticle injection.** Comparing SLC2A1 staining intensity in the syncytiotrophoblast apical (closed arrow) and basal (open arrow) membranes, in non-contemporary untreated controls, intensity was greater in the basal membrane. At 48 h and 10 d following placental *IGF1* nanoparticle treatment, staining intensity of SLC2A1 was similar between the basal and apical membranes. n = 3 placentas per group. Scale bar = 50 µm
